# Emergent microscale gradients give rise to metabolic cross-feeding and antibiotic tolerance in clonal bacterial populations

**DOI:** 10.1101/534149

**Authors:** Alma Dal Co, Simon van Vliet, Martin Ackermann

## Abstract

Bacteria often live in spatially structured groups such as biofilms. In these groups, cells can collectively generate gradients through the uptake and release of compounds. In turn, individual cells adapt their activities to the environment shaped by the whole group. Here we studied how these processes can generate phenotypic variation in clonal populations and how this variation contributes to the resilience of the population to antibiotics. We grew two-dimensional populations of *Escherichia coli* in microfluidic chambers where limiting amounts of glucose were supplied from one side. We found that the collective metabolic activity of cells created microscale gradients where nutrient concentration varied over a few cell lengths. As a result, growth rates and gene expression levels varied strongly between neighbouring cells. Furthermore, we found evidence for a metabolic cross-feeding interaction between glucose fermenting and acetate respiring subpopulations. Finally, we found that subpopulations of cells were able to survive an antibiotic pulse that was lethal in well-mixed conditions, likely due to the presence of a slow growing subpopulation. Our work shows that emergent metabolic gradients can have important consequences for the functionality of bacterial populations as they create opportunities for metabolic interactions and increase the populations’ tolerance to environmental stressors.

## Introduction

Many bacteria live in spatially structured groups where they experience cell densities a hundred to a thousand times higher than in a typical batch-culture [1]. Due to these high densities, cells can modify their environment with their collective metabolic activity [2,3]. Cells living in spatially structured populations can thus create environmental conditions that are not accessible to cells living in isolation and this can have important consequences for the functioning of these populations.

For example, by consuming resources and excreting metabolites cells can generate gradients in the environment [2,3]. In turn, the different microenvironments can induce phenotypic differentiation in the population [4–6]. This differentiation can allow parts of the population to specialize on different tasks, as has been observed in bacterial colonies and biofilms [7–10].

When different subpopulations specialize on complementary metabolic pathways, these subpopulations can engage in cross-feeding interactions [11–14]. Two examples of such metabolic cross-feeding have recently been observed in *Escherichia coli* and in *Bacillus subtilis*. In *E. coli*, cells in the interior of a three-dimensional colony, close to the nutrient source, ferment glucose to acetate, which diffuses to the surface of the colony where it is consumed by a second subpopulation [11,13]. Likewise, in *B. subtilis* it was found that cells in the interior of a two-dimensional biofilm produce ammonium, which is consumed by a second subpopulation on the periphery of the colony [15].

In both these examples metabolic specialization was observed in large populations (of 100’000 cells or more) and the different subpopulations were separated by distances that were much larger than a typical cell length [11,15]. However, in nature bacteria regularly live in nutrient poor environments where population sizes can be much smaller [16]. This raises the question how relevant these metabolic interactions are in natural environments.

Here we hypothesized that phenotypic specialization and metabolic interactions can also occur in small populations and can thus be of general importance in nature. We recently found that, in dense multi-genotype communities, cells are able to create gradients of metabolites on a spatial scale of a few cell lengths, when these metabolites are present in low amounts [17]. In general, the spatial scale over which cells deplete nutrients depends on the density of cells and the concentration of nutrients in the external environment. We thus hypothesized that nutrient gradients could also arise in small clonal populations of densely packed cells growing in nutrient poor environments. As a result, individuals could specialize on different metabolic activities at a very local scale and metabolic cross-feeding could occur even in small populations.

To test our hypothesis, we studied the growth of *E. coli* cells inside microfluidic chambers which open on one side into a flow channel where we supplied low amounts of glucose. In these chambers, cells form two-dimensional populations of about 1000 cells with cell densities comparable to those observed in dense biofilms. While natural biofilms are typically three-dimensional, essential features of life in structured populations can also be observed in two-dimensional populations. This pertains especially to the formation of gradients: typically, nutrients enter a three-dimensional population either from the substrate on which cells grow or from the surrounding liquid environment; in both cases, nutrient gradients are created predominantly along a single dimension, like in our chambers. Therefore, we expect qualitatively similar gradients in two- and three-dimensional populations.

Our microfluidic chambers have multiple advantages: unlike in biofilms, we can easily quantify the properties of individual cells inside the population; furthermore, we can rapidly change the external environment in a controlled way. Our system can thus be used as a model for natural habitats where small populations grow under rapidly changing environmental conditions. Such conditions are for example likely to be found in fluid-filled porous environments, such as the soil [16], or in host compartments such as the lungs in cystic fibrosis patients [18].

We hypothesized that cells in our microfluidic chambers would engage in a glucose-acetate cross-feeding interaction. When glucose is abundant, *E. coli* cells respire glucose only in part, and ferment the rest to acetate [19,20]. In batch cultures glucose and acetate are consumed sequentially: acetate accumulates while cells ferment glucose and is only consumed after glucose becomes depleted [19]. However, in spatially structured populations such as colonies, two different subpopulations are able to metabolize glucose and acetate synchronously [11]. According to our hypothesis, such glucose-acetate cross-feeding could even occur in small populations when the external glucose concentration is low: the cells closest to the glucose source would rapidly consume all available glucose and excrete acetate, while cells only a few cell lengths away would be deprived of glucose and consume the excreted acetate.

We furthermore hypothesized that gradients created by the combined metabolic activity of the population, could provide resilience to changing environmental conditions. As cells adapt their phenotypes to the local conditions in the gradients, phenotypically distinct subpopulations are generated and some of these subpopulations could be more tolerant to stressful environments. For example, biofilms often show tolerance to antibiotic treatments that are lethal to cells growing in batch culture [21]. Tolerance of biofilms to these treatments is partly explained by the formation of slow or non-growing subpopulations [22–24]. We expected that small populations could show similar tolerance to antibiotic treatments whenever strong local gradients in nutrients create subpopulations with high phenotypic variation.

## Results and Discussion

### Cells generate metabolite gradients on micrometre scale

We grew cells inside microfluidic chambers of 40μm wide and 60μm deep, which are closed on three sides and open on one side into a flow channel (Fig 1A). In the microfluidic chambers, which are 0.76μm high, cells grow in two-dimensional monolayers (Fig 1B). In the flow channel we continuously pumped fresh media (M9 minimal medium with 800μM of glucose as the only growth limiting nutrient). In preliminary experiments we established that this concentration is high enough to allow for maximal growth rates close to the chamber opening but low enough to be depleted towards the chamber back. We developed two techniques to automatically quantify the growth of cells: 1) a method quantifying single cell elongation rates using single cell segmentation and tracking and 2) a method quantifying average population growth using optical flow (Fig 1B and Methods).

**Figure 1.**
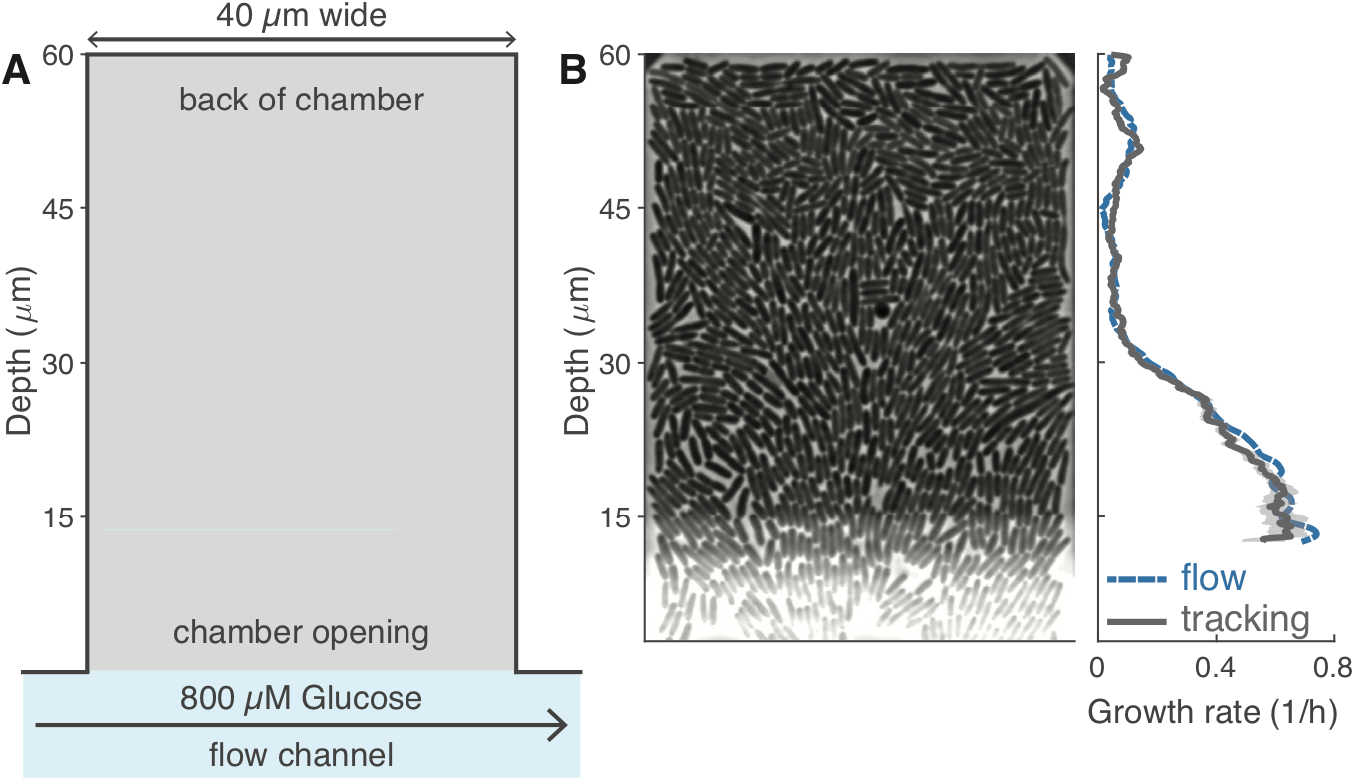
Two-dimensional microfluidic chambers allow for single cell measurements in dense populations. **A)** Cells were grown in chambers of 40*μ*m by 60*μ*m that open on one side into the flow channels. The chambers have a height of 0.76μm forcing cells to grow in a monolayer; the flow channel has a height of 23μm. **B)** Phase contrast image of a single chamber (**left**) with corresponding growth rates as function of depth (**right**). Average growth rates decrease towards the back of the chamber and were estimated using an optical-flow (blue) and cell-tracking (grey) based method. Growth rates were averaged over a one-hour time window, along the width of the chamber, and over a moving window with a depth of 6*μ*m (flow) or 2*μ*m (tracking). Shaded area indicates 95% confidence interval. Image brightness has been adjusted. The phase contrast image shows a band (halo) of higher brightness near the chamber opening: this is an imaging artifact caused by the proximity of the flow channel.

We found that growth rates were maximal close to the chamber opening and decreased (but did not become zero) towards the chamber’s back (Fig 1B, 2B). This suggests that cells collectively form a gradient in glucose along the depth of the chamber and grow slower as glucose becomes depleted. These gradients are created on a micrometre scale: growth rates have decreased by half within 25μm of the chamber opening, a distance that corresponds to only ten cell lengths.

To further quantify the behaviour of cells along the glucose gradient, we measured the expression level of *ptsG*, a high affinity glucose importer that is expressed when cells are exposed to glucose concentrations that are so low that they limit cellular growth rates [25–27]. *ptsG* expression started at depths greater than 30μm and reached maximal expression levels at a depth of 40μm, suggesting that cells in these regions are limited by glucose (Fig 2AB). Near the back of the chamber, *ptsG* expression was almost completely absent, suggesting that glucose is fully depleted in this region (Fig 2AB).

**Figure 2.**
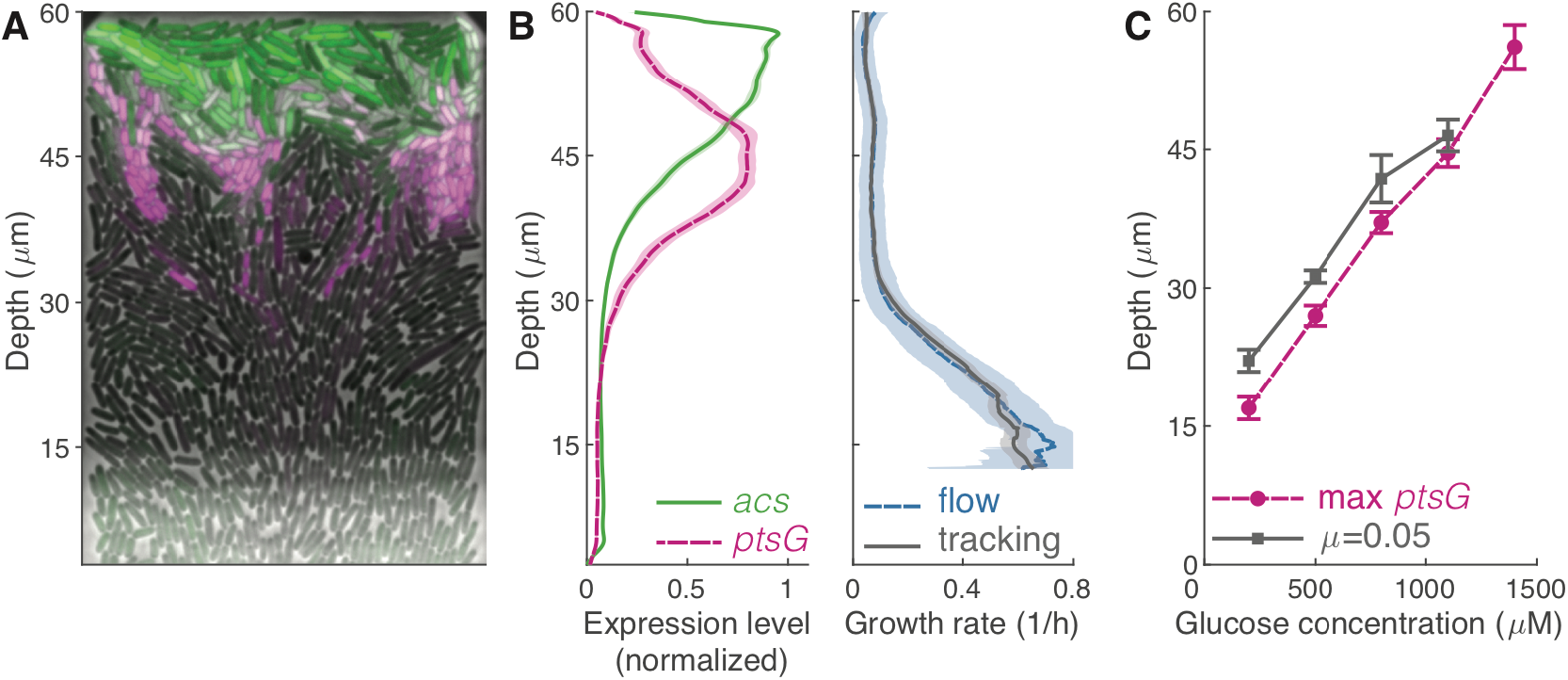
Phenotypic variation arises from nutrient gradients. **A)** False color image of single chamber. Cells limited by glucose express *ptsG* (magenta), cells consuming acetate express *acs* (green). Image brightness and contrast have been adjusted for each color channel separately. **B)** Average gene expression profiles (**left**) and growth rates (**right**) of cells when 800*μ*M of glucose is supplied in the flow channel. Gene expression levels were estimated as the normalized fluorescence intensity of an RFP (*ptsG*) and GFP (*acs*) transcriptional reporter located on the same plasmid. Expression levels were averaged over a moving region with a depth of 2*μ*m and normalized by their maximal values. Profiles show average of three biological replicates (28 chambers), shaded areas indicate 95% confidence intervals. **C)** Depth of gradient measured as the depth at which *ptsG* reaches maximum expression level (magenta circles) or where growth rates reach a value of 0.05 1/h (grey squares) as function of glucose concentration in the flow channel. Data shows average value of ten chambers (five for 1400*μ*M) from one biological replicate, bars indicate 95% confidence intervals

To confirm that the gradients in growth rate and *ptsG* expression are caused by the depletion of glucose, we repeated our experiments while varying the glucose concentration supplied in the flow channel. We observed that both the depth at which growth rate was low (<0.05) and at which *ptsG* expression peaked moved further to the back of the chamber as we increased the glucose concentration (Fig 2C). This data shows that the consumption of glucose by the cells in our chamber depletes glucose to such low concentrations that it leads to a decrease in growth rate along the depth of the chamber. This finding is compatible with previous work in batch and chemostat cultures, where the growth rate of cells was found to reach half its maximal value in a range of glucose concentrations between 200nM to 550μM, depending on culture conditions [28].

Together, our data shows that the collective metabolism of cells generates gradients in glucose. In our chambers, cells likely create gradients in several compounds other than glucose, through the consumption of nutrients and the release of metabolites and signalling molecules. As a result, cells at different locations in the chambers vary strongly in their growth rates and gene expression.

### Nutrient gradients induce emergence of glucose-acetate cross-feeding interactions

We hypothesized that cells near the opening of the chamber produce acetate, which is consumed by cells at the back of the chamber where glucose is not available. To test this hypothesis, we measured the expression level of *acs*, a gene required for growth on low concentrations of acetate [19,27,29]. Expression levels of *acs* peak at the very back of the chamber, where *ptsG* expression decreases (Fig 2AB), suggesting that cells in the back are exposed to acetate but not to glucose. This is consistent with our hypothesis that cells in the back of the chamber grow on the acetate produced by cells near the opening.

To verify that cells in the back of the chamber grow on acetate, we compared the growth of a wildtype strain with that of an Δ*acs* mutant strain. *E. coli* primarily uses the Acs enzyme to grow on low acetate concentrations; Δ*acs* mutant cells should thus be severely impaired in their ability to grow on the low amounts of acetate present in the microfluidic chambers [19,29]. We observed that the Δ*acs* mutant cells consistently grew slower than the wildtype cells in the back of the chamber (Fig 3A). The slow growth we still observed for the Δ*acs* mutant in the back of the chambers could be due to small amounts of remaining glucose, or other metabolites excreted by cells near the chamber opening (e.g. succinate [19]). The Δ*acs* mutant could also potentially metabolize acetate using the PKA pathway [19].

**Figure 3.**
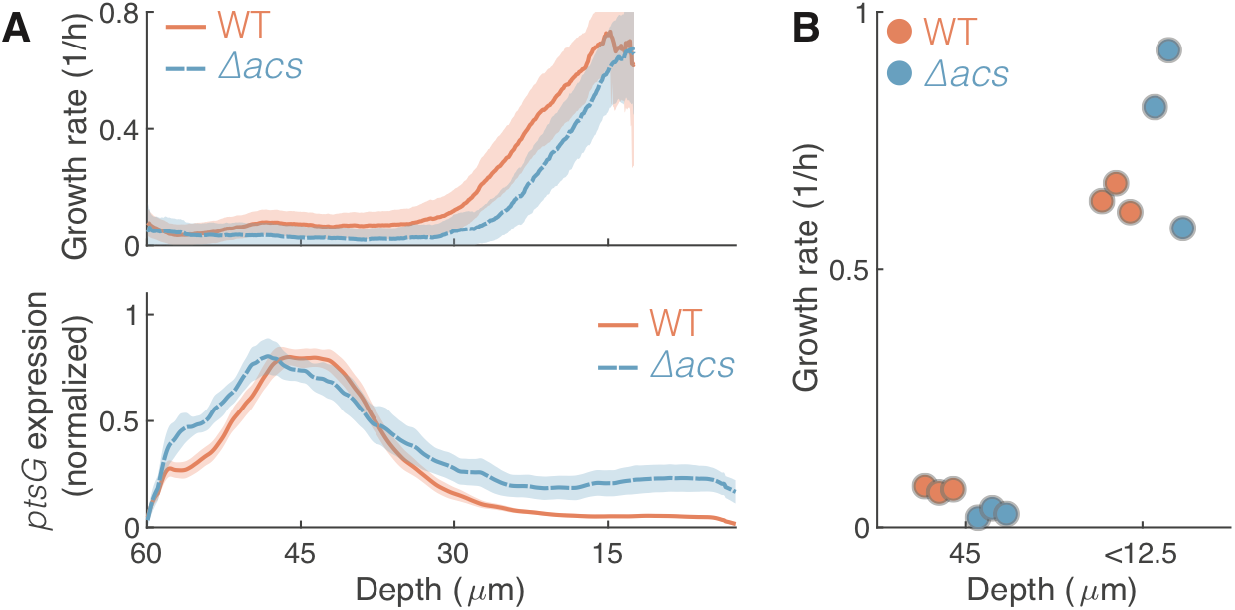
Glucose-Acetate cross-feeding interaction emerges between subpopulations. **A)** Growth rate (**top**) and *ptsG* expression levels (**bottom**) compared between wildtype (WT) and Δ*acs* mutant. Profiles show average of three biological replicates, shaded areas indicate 95% confidence intervals. **B)** The growth rate of wildtype (WT, red) and Δ*acs* mutant (blue) are compared at a depth of 45*μ*m and in the region directly adjacent to the chamber opening (depth < 12.5*μ*m). Each dot corresponds to the average growth rate in a single replicate, with six to ten chambers per replicate at 45*μ*m and one chamber per replicate at 12.5*μ*m. The Δ*acs* mutant strain has a growth defect in the back of the chamber (ANOVA analysis, SI Table 1). In one replicate of the Δ*acs* strain the average growth rate close to the outlet is lower compared to the other two replicates; this is due to the growth curve being shifted by approximately 6*μ*m toward the chamber outlet (see SI Fig. 1).

We quantified the growth defect of the Δ*acs* mutant in the back of the chamber while accounting for the possibility that the two strains have different maximal growth rates. Specifically, we compared the maximal growth rates of cells near the chamber opening (depth < 12.5μm) with the growth rate of cells at a depth of 45μm, where the wildtype is expected to consume acetate (*acs* expression reaches 50% of its maximal level at this depth, Fig. 2B). We performed an ANOVA on log transformed growth rates and found significant effects of strain (*p*<0.001) depth (*p*<0.001) and the interaction between strain and depth (*p*=0.001 3B, SI Table 1). The latter indicates that the wildtype cells have a smaller growth decrease between the chamber opening and the chamber’s back relative to the Δ*acs* mutant. This is consistent with the hypothesis that only wildtype cells consume acetate in the back of the chamber. Our data thus suggest that a cross-feeding interaction occurs between glucose fermenting cells near the opening of the chamber and acetate respiring cells in the back of the chamber.

Glucose-acetate cross-feeding was reported previously in *E. coli* colonies [11,13]. Cole *et al.* observed that above a certain size of the colony two large subpopulations emerge, one fermenting glucose and the other respiring acetate, which are separated by a large non-growing population [11]. Here we found that cells growing within a few cell lengths from each other can differ in their metabolic activity as much as these two subpopulations. Specifically, cells with maximal *ptsG* expression levels are located only five cell lengths (11μm) from cells with maximal *acs* expression levels and the subpopulations expressing *acs* or *ptsG* are both found over a region of only 6 cell lengths in depth (14μm, quantified as the region over which expression levels are above 50% of their maximal value).

Our study complements previous work on metabolic cross-feeding in spatially structured bacterial populations. Specifically, the study by Cole *et al.* used much higher glucose concentrations (14mM versus 0.8mM in our study); as the length scale of the gradient scales with the external glucose concentration, we observed gradients and cross-feeding on much smaller scales. Second, in the previous study, glucose and oxygen entered the colony from opposing sites generating an internal non-growing population separating the glucose and acetate consuming subpopulations; in our system oxygen enters both with the medium (along with the glucose) and through all surfaces of the chamber (the microfluidic devices are fabricated from a material that is permeable to oxygen [30]) and the glucose and acetate consuming cells are located directly adjacent to each other.

Our work, combined with previous studies, shows that cross-feeding interactions between glucose fermenting cells and acetate respiring cells form both in large colonies of billions of cells in nutrient rich environments and in small populations of just a thousand cells in nutrient poor environments. This suggests that glucose-acetate cross-feeding is a robust feature of spatially structured *E. coli* populations. More generally, we can expect cross-feeding interactions to emerge in spatial populations of other bacterial species whenever energetically rich intermediates are released by part of the population [15].

### Cells in structured environments are tolerant to antibiotic exposure

We now turn to our hypothesis that cells in structured environments are more resilient to environmental stressors due to the presence of slow or non-growing subpopulation. We exposed the populations in the microfludic chambers to 50μg/ml streptomycin for three hours. Both before and during antibiotic exposure we supplied low concentrations of glucose (800μM) in the flow channel to allow for the formation of phenotypically diverse subpopulations. After antibiotic treatment we switched to high concentrations of glucose (10mM) to test whether cells in the back of the chamber survived the antibiotic treatment and could grow.

We observed that spatially structured populations were tolerant to antibiotic exposure. While in batch cultures a three-hour pulse of 50μg/ml streptomycin led to more than a two-million-fold reduction in cell numbers, we observed that in all 38 chambers (with each about 1000 cells) parts of the populations were able to survive the antibiotic pulse (Fig 4A).

**Figure 4.**
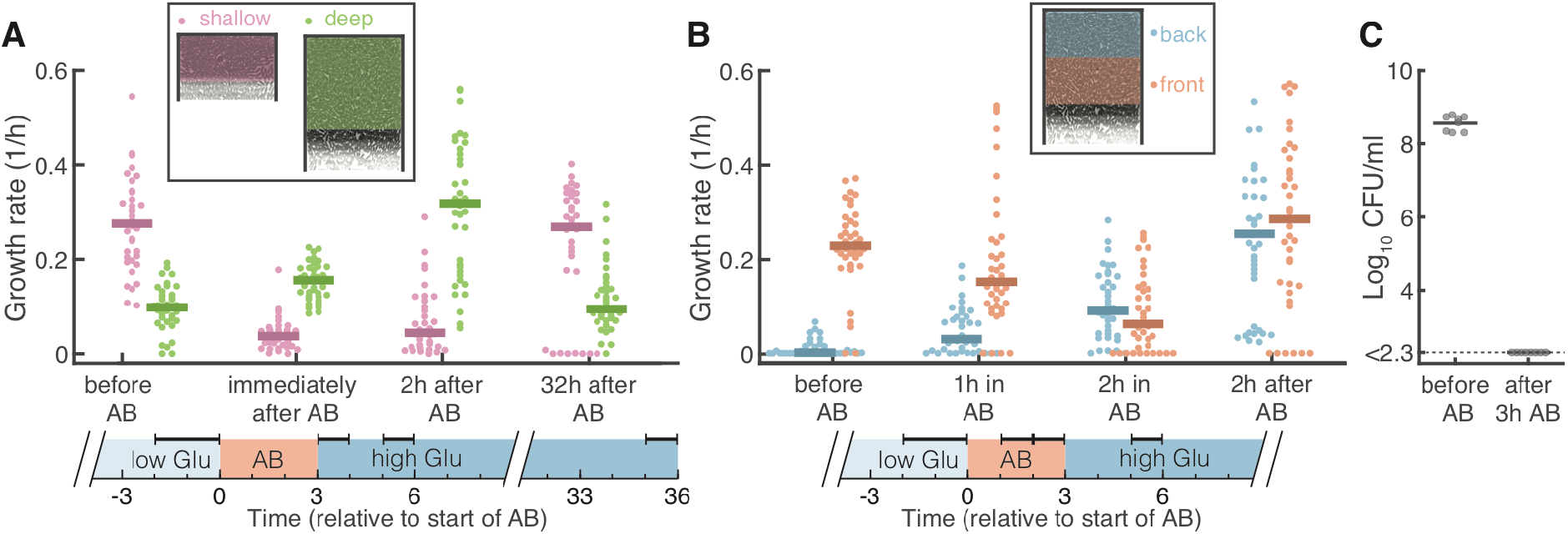
Spatially structured populations are tolerant to antibiotics. Cells were first grown on low glucose medium for nine hours, exposed to three hours of 50*μ*g/ml streptomycin (AB), and finally grown in high glucose medium for 35 hours. **A)** Cells in 60*μ*m deep chambers (green) cope better with antibiotic treatment than cells in 30*μ*m shallow chambers (pink). In shallow chambers population growth rates decrease strongly during antibiotic treatments and take a long time to recover (eight chambers have zero growth even after 32 hour), while in deep chambers growth rates remain high during the entire treatment. Horizontal bars on top of timeline indicate the analyzed time periods. **B)** Cells in the back of the chamber (blue) tolerate antibiotics better than cells in the front (red). During antibiotic treatment, the average growth rates in the front of the chamber decreases, while that in the back of the chamber increases. Two hours after treatment, the average growth rate in the front of the chamber is high again as the non-growing population is fully replaced by growing cells coming from the back (see SI Movie 1). Horizontal bars on top of timeline indicate the analyzed time periods. **A,B)** Each dot shows the average population growth rate inside the colored region shown in the schematics. Negative growth rates were set to 0. Bars show median values. Growth rates were averaged over the time windows indicated by the horizontal bars at the top of the timelines. **C)** Cells in batch cultures cannot survive antibiotic treatment. Colony forming units (CFU) per ml are shown for 8 replicate batch cultures just before and three hours after addition of 50*μ*g/ml streptomycin. After three hours of antibiotic exposure all 8 cultures had a CFU count below the detection limit (of 200 CFU/ml), corresponding to a decrease in CFU counts by at least a factor 2 · 10^6^.

We hypothesized that the populations in the microfluidic chambers survived antibiotic treatment due to the increased tolerance of the slow growing subpopulation of cells in the back of the chamber. Visual inspection of the chambers suggested indeed that most cell death happened near the opening of the chamber, while cells in the back could survive the antibiotic treatment (SI Movie 1). To test this idea, we compared the effect of antibiotic treatment in our standard *deep chambers* (depth=60μm) with that in *shallow chambers* (depth=30μm).

We observed that populations in deep chambers could cope much better with the antibiotic treatment than populations in shallow chambers. Before antibiotic treatment, the growth rate of cells in the shallow chambers was similar to those observed in the front 30μm of the deep chambers but the slow and non-growing subpopulations found in the back of the deep chambers was not present in the shallow chambers (SI Fig. 3). During antibiotic exposure population growth rates decreased in both deep and shallow chambers, however they recovered much faster in the deep chambers: two hours after removing the antibiotic the average growth rate in the deep chambers was as high as before the antibiotic treatment, while the average growth rate in the shallow chamber was much lower (Fig 4A). This lower growth rate was largely due to a difference in cell survival: whereas in deep chambers a large fraction of cells survived (SI Movie 1), in shallow chambers often only a single or very few cells could resume growth after the antibiotic treatment (SI Movie 2). The low survival in the shallow chamber also explains why eight out of 37 populations in the shallow chambers went extinct, while all 38 populations in the deep chambers survived antibiotic exposure.

To further test whether cells in the back of the chambers were indeed the main source of population survival, we compared the average growth rates between the front and the back of the chamber. Before antibiotic exposure cells in the back on average grew slower than cells in the front as a result of the glucose gradient (Fig 4B). However, during antibiotic treatment cells in the back started growing faster while the ones in the front rapidly stopped growing (Fig 4B). Why did the cells in the back grow faster during the antibiotic treatment than before? Most likely, as cells in the front of the chamber died, nutrients became available for cells in the back, allowing them to grow faster. Directly after antibiotic exposure, these cells grew even faster as a result of the high glucose concentration we supplied in the flow channel. While most cells in the back had survived the antibiotic treatment, most cells at the front were not growing (and presumably dead), thus causing the low average growth rate at the front (Fig 4B, SI Movie 1). Two hours after the end of the antibiotic pulse, most non-growing cells at the front were pushed out of the chamber and replaced by rapidly growing cells from the back. As a result, the difference between the front and back of the chamber disappeared (Fig 4B).

Together our data supports the hypothesis that many fast-growing cells near the chamber opening die, while the slow growing cells in the back of the chamber are able to survive the antibiotic treatment. This finding is consistent with a number of previous studies that found a positive correlation between slow growth and antibiotic survival [22–24,31]. While we cannot exclude that other factors (e.g. gradients in antibiotic concentration) also contribute to the survival of cells at the back of the chambers, their low growth rate, which result from the emergent glucose gradient, could play a decisive role in their survival.

## Conclusion

In dense, spatially structured environments, cells can collectively change their environment and create strong gradients in metabolites. Generally, whenever cellular densities are high and nutrient concentrations are low, steep gradients can occur locally, on length scales of only a few cell lengths. As cells adapt to their local conditions, these gradients give rise to phenotypically distinct subpopulations that specialize at a very local scale on different metabolic tasks and that can engage in metabolic cross-feeding interactions.

Although our study focused on *E. coli*, we expect our findings to be of direct relevance to natural populations of many microbial species, as often the natural environments of bacteria and other microorganisms are characterized by low nutrient availability and dense spatially structured populations. Microscale phenotypic variation can thus be an important aspect of natural microbial populations. The experimental and analysis techniques developed in this work provide tools to investigate in details how bacteria behave in dense spatial populations and how they achieve collective functionality.

Microscale phenotypic variation can have important consequences for the functionality of bacterial populations as it promotes metabolic interactions between distinct subpopulations and tolerance to environmental stressors. Living together in structured environments thus allows cells to achieve functionality collectively that they cannot achieve alone.

## Methods

### Strains and plasmids

All experiments were done with *E. coli* MG1655 (WT) or *E. coli* MG1655 *acs∷frt* (Δ*acs* mutant) carrying the low copy number pSV66-*acs-gfp-ptsG-rfp* dual transcriptional reporter. This plasmid was constructed from pSV66-*rpsM-gfp-rpsM-rfp* by replacing the promoter sequences upstream of *rfp* and *gfp* using a one-step Gibson assembly [32]. The following 4 fragments were amplified using the Q5 high fidelity polymerase (NEB) and combined using Gibson assembly (NEB): 1) *acs* promoter, amplified from plasmid pUA66-*acs-gfp* [14] using primer pGFP-fw and pGFP-rv; 2) *ptsG* promoter, amplified from pUA66-*ptsG-gfp* [33] using primers pRFP-fw and pRFP-rv; 3) pSV66 *gfp-rfp* region, amplified from pSV66-*rpsM-gfp-rpsM-rfp* using primers GFP_vec-fw and RFP_vec-rv; 4) pSV66 backbone region, amplified from pSV66-*rpsM-gfp-rpsM-rfp* using primers GFP_vec-rv and RFP_vec-fw (see SI Table 2 for primer sequences). The sequences of the promoter regions were verified with Sanger sequencing. Kanamycin was added to all growth media to select for plasmid maintenance. However, even in the absence of kanamycin, we expect plasmid loss to be minimal during the duration of our experiment because the reporter plasmid carries the pSC101 replication origin which has very low copy number variation [34]. The *E. coli* MG1655 *acs∷frt* was constructed from MG1655 *acs∷kanR* obtained from the Keio collection [35], by removal of the kana-mycin cassette using Flp-FRT recombination.

### Media and growth conditions

Cells were grown in M9 medium (47.76 mM Na_2_HPO_4_, 22.04 mM KH_2_PO_4_, 8.56 mM NaCl and 18.69 mM NH_4_Cl) supplemented with 1mM MgSO_4_ and 0.1 mM CaCl_2_ (all from Sigma-Aldrich). Glucose was added to the medium to a final concentration of 10mM (high glucose medium), 800μM (low glucose medium), or as specified in the figure captions. All media was supplemented with 50μg/ml kanamycin and 0.1% of Tween-20 (Polysorbate-20, Sigma-Aldrich) to reduce sticking of cells to the sides of microfluidic devices. Overnight cultures were started from a single colony from a LB agar plate and grown in M9 medium supplemented with 10mM glucose and 5% LB. All cultures were grown at 37°C in a shaking incubator. For the experiments on antibiotic tolerance in the microfluidic chambers 50μg/ml of streptomycin was added to the low glucose medium.

We used two different batches of M9 growth media (same manufacturer and article number, but different lot number) and observed some quantitative differences between experiments done with the different batches: growth and expression profiles were shifted towards the back of the chamber in one batch compared to the other. We observed qualitatively the same results with both batches of media and all our conclusions are robust to differences between the batches. To account for the quantitative difference between the two batches, we never made direct comparison between data obtained from experiments done with different batches: specifically, we used batch one to measure how the gradient changes with glucose concentration and how the cells respond to antibiotics (Fig 2C, 4AB) and batch two for all other experiments.

### Antibiotic tolerance in well-mixed conditions

Eight independent cultures were started from separate colonies from a LB agar plate and grown overnight in LB medium supplemented with 50μg/ml kanamycin. The next day cultures were diluted to a final optical density at 600nm (OD600) of 0.1 into 1.5 ml of fresh LB medium (supplemented with 50μg/ml kanamycin) in a 24-well plate and grown for three hours to mid-exponential phase. From each culture, a 20ul sample was taken and a dilution series was spot-plated on LB agar plates to obtain the cell density before antibiotic exposures, measured as colony forming units (CFUs). Subsequently, streptomycin was added to a final concentration of 50μg/ml (MIC < 25μg/ml) and the cultures were grown for another 3 hours before being sampled again to obtain the cells densities after antibiotic exposure. All cultures were grown at 37°C in a shaking incubator.

Right before antibiotic exposure cell densities were 4.0 ± 1.2 · 10^8^ CFU/ml (mean ± 95% confidence interval, *n*=8). After antibiotic exposure we had no detectable CFUs in any replicate. As our detection limit was 200 CFU/ml. This implies that the antibiotic exposure caused at least a two-million-fold decrease in cell densities.

### Microfluidic devices

Molds for the microfluidic devices were constructed using a two-layer photolithography process using SU8 photoresist on Silicon wafers. Microfluidic devices were made by pouring polydimethylsiloxane (PDMS, Sylgard 184) on the SU8 molds, after which air bubbles were removed using a desiccator before baking the devices at 80°C for one hour. Subsequently, the PDMS devices were bound to microscope cover glass slides by treating them with oxygen plasma (PDC-32G-2 Plasma Cleaner, Harrik Plasma, New York, USA), and leaving them on a heated plate at 100°C for one minute.

The device consists of a long (≈2cm) flow channel of 100μm wide and 23μm high which connects to chambers that are 0.76μm high, 40μm wide and 30 or 60μm deep. The small height of the chambers ensures that cells grow in a monolayer. In preliminary experiments we observed that the height of the chambers is of critical importance: when chambers are too high (>0.8μm), cells are lost from the chambers easily and they cannot form a densely packed layer; when chambers are too low (<0.7μm), cell growth is impaired. We chose a height of 0.76μm as this is the lowest value that allows for normal cell growth. We expect some variation in height within each chamber because cells exert pressure and can deform the elastic PDMS material.

### Microscopy

Time-lapse microscopy was done using a fully-automated Olympus IX81 inverted microscope controlled with the CellSens software (Olympus). Imaging was done with a 100X NA1.3 oil phase objective (Olympus) and an ORCA-flash 4.0 v2 sCMOS camera (Hamamatsu). Fluorescent imaging was done with the Chroma N49002 (GFP) and N49008 (RFP) filters and X-Cite120 120-Watt high pressure metal halide arc lamp (Lumen Dynamics). The Olympus Z-drift compensation system was used to maintain focus. A microscope incubator (Life imaging services) maintained the sample at 37°C.

### Microfluidic Experiments

The microfluidic devices were wetted with culture medium using a pipette to facilitate subsequent cell loading. An overnight culture of cells was concentrated by centrifugation and loaded into each flow channels by pipette and cells were pushed into the side chambers. Subsequently the inlets of the flow channels were connected via tubing (Microbore Tygon S54HL, ID 0.76 mm, OD 2.29 mm, Fisher Scientific) to 50ml syringes and media was flown continuously at 0.5ml/h using syringe pumps (NE-300, NewEra Pump Systems).

#### Response to nutrient gradients

Cells were first grown in high glucose medium (10mM) until they had filled the chambers (≈18 hours). Subsequently the medium was switched to the low glucose medium (800μM) for nine hours. In preliminary experiments we established that it takes about three to four hours for cells to form and adapt to the nutrient gradients. To ensure that all measurements were taken at steady state we thus started imaging the chambers six hours after switching to low glucose medium.

We imaged the population with two regimes: for the first hour we imaged in phase contrast, taking an image every one minute and 45 seconds; for the next two hours we imaged the population in phase contrast (to measure biomass), GFP (*acs* expression), and RFP (*ptsG* expression) taking an image every six minutes.

We used this strategy because accurate determination of growth rates (using either optical flow or single cell tracking techniques) requires high frequency imaging, which is not compatible with the time required for the acquisition of fluorescent images.

#### Response to antibiotic pulse

Cells were first grown in high glucose medium (10mM) until they had filled the chambers (≈18 hours). Subsequently the medium was switched to the low glucose medium (800μM) for nine hours, to establish the gradient in growth rates. Finally, cells were submitted to antibiotic treatment and recovery, applying 3 hours of low glucose medium with 50μg/ml streptomycin, followed by 35 hours of high glucose medium without streptomycin. We used high glucose medium after the streptomycin pulse to ensure that all cells had access to glucose so we could observe growth of surviving cells irrespective of their location in the chamber.

The population was imaged in phase contrast (to measure biomass), GFP (*acs* expression), and RFP (*ptsG* expression) taking an image every six minutes for 40 hours. The imaging was started two hours before the start of the (three hour long) antibiotic pulse and continued for 35 hours after the end of the pulse. To compare the growth rate profiles in 30 and 60μm deep chambers we furthermore imaged a subset of the data at high frequencies (taking an image every one minute and 45 seconds) in phase contrast only for one hour starting three hours before the antibiotic pulse.

### Image analysis

Time-lapse movies where analysed using the custom build Vanellus image analysis software (Daan Kiviet, [17]), Ilastik [36] and Matlab (version 2016a and newer). Movies were registered to compensate for stage movement and cropped to the region of the growth chambers.

#### Segmentation

Cells were segmented using the phase contrast images and two independent segmentation techniques. We segmented the single cells using Ilastik software based on supervised machine learning. The Ilastik classifier was trained to maximize accuracy in identification of individual cells (i.e. to accurately separate nearby cells). As this technique tends to underestimate the total biomass (part of the cell contour is excluded), we used a second technique to accurately identify total biomass in the chamber. This second technique is optimized to maximize accuracy in detection of biomass, without attempting to separate neighbouring cells. This technique was implemented in custom built Matlab code and uses and algorithm combining edge detection with subsequent filtering on cell size and texture (Hessian eigen values).

#### Tracking

Cell tracking was done using the optical-flow based tracking algorithm described in ref. [17]. In short: cells were tracked by estimating the movement between two subsequent images with optical flow and using this to predict the location of each cell in subsequent frames.

#### Fluorescent images

Fluorescent images were deconvolved using the Lucy-Richardson method and back-ground corrected as 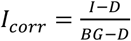, where *I* is the uncorrected background intensity, *D* is the median pixel value of an image taken with closed shutter (dark count), and *BG* is the background intensity measured in the flow channel directly adjacent to the chamber exit.

### Growth rate calculation

#### Population average growth rate

The average population growth rate can be estimated as follows: Consider a region in the chamber of width Δ*y* centered at depth *y* (the y axis is oriented from the chamber opening to the chamber’s back), and let *B*(*t*, *y*) be the biomass in this region at time *t*. During a time period Δ*t* the biomass *B*(*t*, *y*) varies due to growth and movement of biomass in and out of the region. The variation due to growth is given by *μ*(*t*, *y*) · *B*(*t*, *y*) · Δ*t*, where *μ*(*t*, *y*) is the average population growth rate; the variation due to movement is the balance between movement of biomass in and out of the region, thus if Φ(*t*, *y*) is the velocity field this is given by: 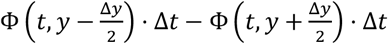. The total change in biomass Δ*B*(*t*, *y*) is thus given by:

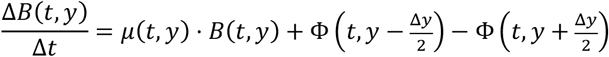

This equation can be used to calculate the growth rate as:

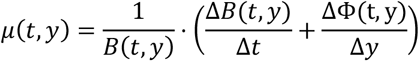

We estimated *B*(*t*, *y*) from the segmented images as the number of pixels occupied by cells, and we estimated Φ(t, y) using Farneback optical flow algorithm applied to the phase contrast images [37]. All quantities are calculated over a time window Δ*t* around t (typically 1h) and over a spatial region Δ*y* around y: *B*(*t*, *y*) is the biomass averaged over Δ*t* and 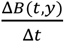 is the average slope of the linear regression of *B*(*t*, *y*) with *t*, for *t* within Δ*t*; 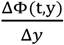 is the slope of the linear regression of Φ(t, y) with y, for *y* within Δ*y*. The main source of error in this estimation of growth rates is the velocity field Φ, which sometimes wrongly detects movement between two frames. To automatically correct for this, we excluded the four time points (10%) with the worst quality before averaging 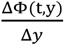 over the remaining time points. The quality of each time point was automatically assessed as the mean squared error of the linear regression averaged over all *y* for a given time point.

The velocity field cannot be correctly estimated close to the opening of the chamber because of the presence of a halo in the phase contrast images (this imaging artefact is well visible around ~12.5μm from the chamber opening and is caused by the proximity of the flow channel). The growth rate cannot be estimated in this halo region using the formula above. As the halo region is small, we can still estimate the growth profile along most of the chamber depth and particularly how growth decreases with depth. The growth profiles are automatically cutoff at the depth where they reach their maximal value to exclude regions where growth rate cannot be estimated well with this method.

#### Single cell growth rates

Single cell growth rates were measured as the elongation rate of cells: *l*(*t*) = *l*(0) · *e*^*μ*·*t*^; *μ* was estimated as the slope of the linear regression of the log-transformed cell length over a moving time window of 7 points (12 minutes). Thus, only cells for which length measurements were available for at least 7 time points were included in the analysis. Both cell segmentation and tracking were fully automated with no manual correction applied at any stage. To automatically exclude tracking mistakes, we screened for cells whose length displayed large fluctuation in time. This was done by calculating the reduced Chi-squared value for each regression as 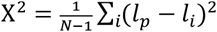, where N is the number of time points over which the regression is done, *l*_*p*_ is the length predicted by the linear regression and *l*_*i*_ is the measured cell length. Visual inspection of a subset of the data suggested that excluding trajectories with a Χ^2^ > 4 · 10^−4^ resulted in reliable estimates of single cell growth rates.

### Single cell growth rates near the chamber opening

Within 10 *μ*m of the chamber opening the automatic procedure described above is no longer able to measure single cell growth rates. The rapid movement of cells in this region complicates automatic tracking, while the phase contrast artefact caused by the proximity of the flow channel affects both the automatic segmentation and tracking quality. To still obtain growth rates in this region for both the wild type and Δ*acs* mutant populations, we selected a subset of our data (one chamber per replicate) based on the quality of the image segmentation. We subsequently manually corrected the segmentation and tracking of cells in the region up to 20*μ*m from the opening for twelve frames (17 minutes). Afterwards, the single cells growth rates were obtained as described above.

### Statistical treatment of data

#### Data exclusion

We only included chambers that were fully packed with cells during the full observation window, based on visual inspection. Chambers where part of the population was lost at any time (large groups of cells leaving the chamber likely because of pressure fluctuations) were excluded from the analysis.

#### Spatial averaging

Our system is quasi-1D: there is no systematic variation along the width of the chamber. We thus averaged all quantities along the chamber width to obtain 1D profiles of phenotype versus depth in the chamber. To obtain smooth profiles along the depth of the chamber we used moving averages. Specifically, population average growth rates (optical flow based) were calculated over a 6μm (91 pixel) window, while single cell growth rates and fluorescent profiles were averaged over a 2μm (31 pixel) window.

#### Time averaging

All gradient measurements where averaged over 1h. Growth rates and gene expression level measurements were taken in two non-overlapping but directly adjacent time windows. As the growth rate and gene expression profiles are approximately constant during the experiment we can superpose these two measurements despite the small time offset. For the analysis of the response to antibiotics, growth rates before antibiotic treatment were averaged over a time window of two hours (for one replicate only 6 minutes of data were recorded before the switch, in this case averaging was done only over this interval); growth rate during and after antibiotic treatment were averaged over a time window of one hour.

#### Replicates

For the gradient measurements we imaged three flow channels with nine to ten chambers each (28 chambers in total) for the wildtype and three flow channels with six to nine chambers (22 chambers in total) for the Δ*acs* mutant. Each flow channel was inoculated with a different overnight culture and was considered to be an independent biological replicate. We treated chambers within the same flow channel as technical replicates. For the antibiotics we measured 38 deep and 37 shallow chambers, both from the same four independent flow channels.

#### Chamber averaging

For the data shown in Figures 1-3 all chambers of a given flow channel were first averaged together. Simple averages were used for the optical flow based population growth rate and fluorescent measurements; weighted averages were used for the single cell growth rates, with weights corresponding to the number of cells measured in each chamber at a given depth (chambers with less than 20 cells at a given depth were excluded from the analysis).

### Data availability

The data and code required to reproduce the figures and conclusions are available on the Scholars Portal Dataverse repository: https://doi.org/10.5683/SP2/CBVYXB [38].

## Acknowledgements

We thank Daan Kiviet for providing the Vanellus image analysis software and the molds for the microfluidic devices, Jana Fahrion for performing initial exploratory experiments, Jasmin Annaheim help with the image analysis, and Susan Schlegel, Glen Dsouza, and the other members from the microbial systems ecology group for helpful feedback and discussions.

This work was supported by the Swiss National Science foundation (grants no. 31003A_149267 and 31003A_169978 to MA), by an Early Postdoc Mobility fellowship from the Swiss National Science Foundation (grant no. 175123 to SVV), and by ETH Zurich and Eawag.

## Author Contributions

ADC designed the study, collected, analyzed, and interpreted the data, and drafted the manuscript; SVV designed the study, engineered the genetic constructs, collected, analyzed and interpreted the data, and drafted the manuscript; MA designed the study and critically revised the manuscript. All authors gave final approval for publication and agree to be held accountable for the work performed therein.

