## Supplemental Information for "Emergent microscale gradients give rise to metabolic cross-feeding and antibiotic tolerance in clonal bacterial populations"

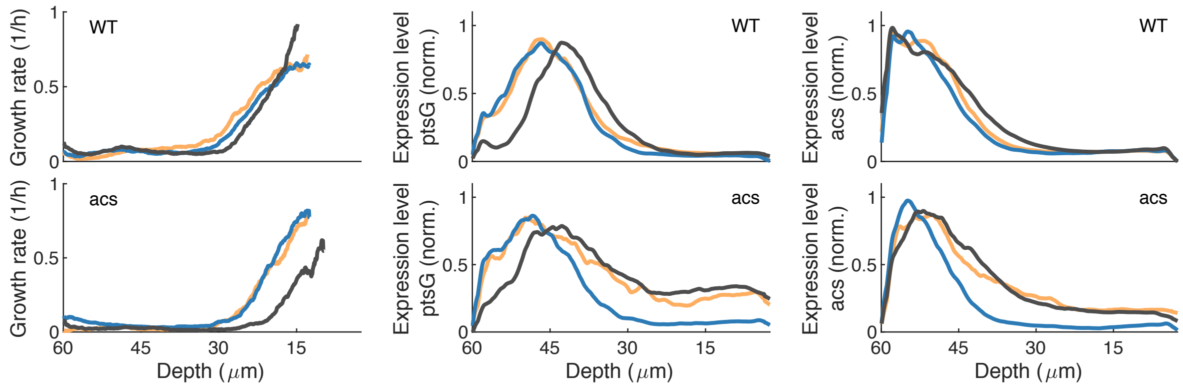


**Supplemental Figure 1.** Nutrient gradients and gene expression are reproducible across replicates. The figure shows the profiles of single cell growth rate (left), normalized ptsG expression (middle), and normalized acs expression (right) for wildtype (WT, top) and acs mutant populations (bottom), each line shows average values within a single replicate (six to ten chambers). Note that overall profiles are very similar between replicates, though some are shifted slightly towards higher or lower depth (e.g. dark gray line *acs* mutant). This is likely due to slight variation in the density of cells, chamber height, or flow pressure that can change how deep glucose can penetrate into the chamber before being depleted.

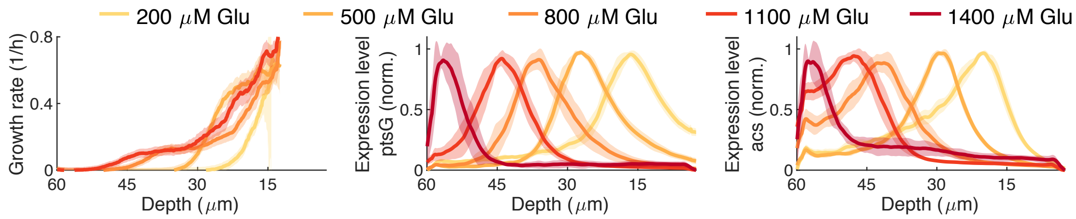


**Supplemental Figure 2.** Depth profiles of *ptsG* expression (left), *acs* expression (middle), and single cell growth rate (right), different colors correspond to different glucose concentration in the flow channel. The peaks of *ptsG* and *acs* expression move to deeper depths, as the glucose concentration in the flow channel is increased. At the same time the growth rates of cells increase in the back of the chamber and a distinct plateau is formed, likely due to growth on acetate in this region. Note that the profiles shown here for 800$\mu$M are slightly different from those shown in Fig. 2 as these experiments were done using different batches of M9 media (see Methods for details). Importantly, we see qualitatively the same behavior between the different media batches. Lines show average over ten chambers (five for 1400$\mu$M) from one biological replicate, no growth rate data was available for 1400$\mu$M glucose.

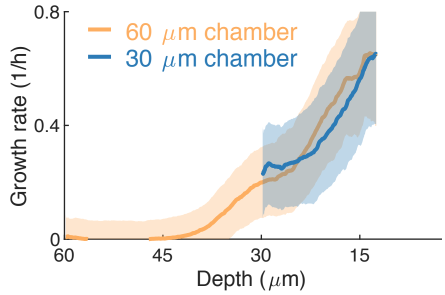


**Supplemental Figure 3.** Depth profile of average single cell growth rate for chambers with a depth of 60$\mu$m (orange, *deep chambers*) and 30$\mu$m (blue, *shallow chambers*). Note that the growth rate of cells in the shallow chambers is similar to the growth rate of cells the front half of the deep chambers, while the slow and non-growing populations observed in the deep chambers are absent from the shallow chambers. Lines show average over two biological replicates (with six to ten chambers each), shaded area indicates 95% confidence interval.

**Supplemental Movie 1.** Response of population growing in *deep chamber* (depth=60$\mu$m) to antibiotic exposure. Cells were grown in 800$\mu$M glucose, exposed for three hours to 50$\mu$g/ml of streptomycin, and then switched to 10mM glucose; time and medium in flow channel are shown in top left corner. Note how many cells near chamber opening stop growing (and presumably die) during the antibiotic pulse, while most cells in the back of the chamber survive and can keep growing. After the end of the antibiotic pulse the growing cells in the back of the chamber rapidly push out all non-growing cells near the chamber opening, thus restoring the full population.

**Supplemental Movie 2.** Response of population growing in *shallow chamber* (depth=30$\mu$m) to antibiotic exposure. Cells were grown in 800$\mu$M glucose, exposed for three hours to 50$\mu$g/ml of streptomycin, and then switched to 10mM glucose; time and medium in flow channel are shown in top left corner. Note how almost all of the cells in the chamber stop growing (and presumably die) during the antibiotic pulse. After the end of the antibiotic pulse only a single cell can resume growth (indicated by arrow) and even after eight hours the majority of the cells in the chamber cannot grow.

**Supplemental Table 1.** Anova table of comparison between growth of wildtype and *acs* mutant strain between front (depth = 12.5$\mu$m) and back (depth = 12.5$\mu$m) of chamber. Growth rates were log transformed.

| **Source** | **Sum of squares** | **Degrees of freedom** | **Mean squares** | **F-statistic** | ***p*** |
|---|---|---|---|---|---|
| **Strain** | 0.55 | 1 | 0.55 | 12.1 | $8 \cdot 10^{-3}$ |
| **Depth** | 22.90 | 1 | 22.90 | 503.1 | $2 \cdot 10^{-8}$ |
| **Strain*Depth** | 1.09 | 1 | 1.09 | 23.9 | $1 \cdot 10^{-3}$ |
| **Error** | 0.36 | 8 | 0.05 | | |
| **Total** | 24.90 | 11 | | | |

**Supplemental Table 2.** Primer sequences used in this study. Underlined parts are tails added to primers to provide homology for Gibson assembly.

| Name | Sequence | Target | Target Source |
|---|---|---|---|
| **pGFP-fw** | 5’ CTG GCA ATT CCG ACG TCT AAG AAA C | pUA66-*acs-gfp* | [1] |
| **pGFP-rv** | 5’ CAA CAA GAA TTG GGA CAA CTC CAG TG | pUA66-*acs-gfp* | [1] |
| **pRFP-fw** | 5’ TAA CAA ACT AGC AAC ACC AGA ACA GGT ATC ACG AGG CCC TTT CG | pUA66-*ptsG-gfp* | [2] |
| **pRFP-rv** | 5’ TGT TCT CCT TGA TCA GCT CGC TCA TAT GTA TAT CTC CTT CTT AAA TCT AGA GGA TCC | pUA66-*ptsG-gfp* | [2] |
| **GFP_vec-fw** | 5’ CAC TGG AGT TGT CCC AAT TCT TGT TG | pSV66-*rpsM-gfp-rpsM-rfp* | [3] |
| **RFP_vec-rv** | 5’ ATG AGC GAG CTG ATC AAG G | pSV66-*rpsM-gfp-rpsM-rfp* | [3] |
| **GFP_vec-rv** | 5’ GTT TCT TAG ACG TCG GAA TTG CCA G | pSV66-*rpsM-gfp-rpsM-rfp* | [3] |
| **RFP_vec-fw** | 5’ CTG TTC TGG TGT TGC TAG TTT G | pSV66-*rpsM-gfp-rpsM-rfp* | [3] |
